# The contribution of hippocampal subfields to the progression of neurodegeneration

**DOI:** 10.1101/2020.05.06.081034

**Authors:** Kichang Kwak, Marc Niethammer, Kelly S. Giovanello, Martin Styner, Eran Dayan, for the Alzheimer’s Disease Neuroimaging Initiative

## Abstract

Mild cognitive impairment (MCI) is often considered the precursor of Alzheimer’s disease. However, MCI is associated with substantially variable progression rates, which are not well understood. Attempts to identify the mechanisms that underlie MCI progression have often focused on the hippocampus, but have mostly overlooked its intricate structure and subdivisions. Here, we utilized deep learning to delineate the contribution of hippocampal subfields to MCI progression using a total sample of 1157 subjects (349 in the training set, 427 in a validation set and 381 in the testing set). We propose a dense convolutional neural network architecture that differentiates stable and progressive MCI based on hippocampal morphometry. The proposed deep learning model predicted MCI progression with an accuracy of 75.85%. A novel implementation of occlusion analysis revealed marked differences in the contribution of hippocampal subfields to the performance of the model, with presubiculum, CA1, subiculum, and molecular layer showing the most central role. Moreover, the analysis reveals that 10.5% of the volume of the hippocampus was redundant in the differentiation between stable and progressive MCI. Our predictive model uncovers pronounced differences in the contribution of hippocampal subfields to the progression of MCI. The results may reflect the sparing of hippocampal structure in individuals with a slower progression of neurodegeneration.

## Introduction

A certain degree of cognitive decline is common and considered a part of the normal aging process. Mild cognitive impairment (MCI) occurs when cognitive decline exceeds what is expected given an individual’s age and education level (1). MCI can be considered a transitional phase in between age-related cognitive decline and Alzheimer’s disease (AD) or other dementias (1). However, MCI is associated with marked etiological heterogeneity (2) and variable progression rates (3). Namely, up to 33% of individuals with MCI convert to AD over 5 years (4), with annual conversion rates of about 7% (5), but others may remain stable or even revert to normal or near-normal cognition levels (6). Identifying prognostic markers that can predict eventual conversion from MCI to AD is of profound clinical interest (7). However, such markers are not available to date. Fundamentally, despite increased interest in recent years (8–10), a mechanistic framework for understanding progression and stability in MCI remains missing.

The neuropathological profile of MCI is complex and multifaceted (11). As MCI is commonly seen as a precursor to AD, many studies have focused on alterations in structures known to be affected by this disease. The most widely studied target in AD is the hippocampus (12, 13), and indeed, multiple studies have reported hippocampal volume loss in MCI relative to controls (14). Studies have also implicated the hippocampus in the progression of MCI (15). Of particular interest, a series of recent studies have utilized deep learning to differentiate progressive and stable MCI (16, 17), or predict individual subjects’ progression from MCI to AD (18) based on whole hippocampus structural features. However, rather than being a homogeneous structure, the hippocampus is complex and heterogeneous (19). The hippocampus is composed of several histologically distinct subfields (19), which are characterized by differential connectivity profiles (20) and subserve different memory processes (21). Thus, a better understanding of MCI progression and stability necessitates a mechanistic framework that takes the structural complexity of the hippocampus into account (22). Whether the different hippocampal subfields contribute differentially to the progression from MCI to AD remains unclear, with mixed and inconsistent findings reported in the literature. Namely, while several studies reported that the CA1 and subiculum are the most central subfields in the progression of MCI (23, 24), other studies suggested that the CA2/3, fimbria, and GC-DG were most central (25, 26).

In the current study, we investigated the contribution of hippocampal subfields to the progression and stability of MCI. To that end, we utilized a deep learning framework, and a large neuroimaging dataset, to account for the expected complexity in the contribution of the different subfields to the progression of MCI. We propose a deep convolutional neural network trained to classify stable and progressive MCI based on hippocampal structural features derived from magnetic resonance imaging (MRI). We then introduce a novel implementation of occlusion analysis to evaluate the relative contribution of each hippocampal subfield to the performance of the predictive model. Moreover, the same analysis allowed us to estimate the cumulative contribution of subfields to MCI stability and the possible existence of redundancy within the associated hippocampal features.

## Results

To delineate the contribution of hippocampal subfields to MCI progression we analyzed neuroimaging data from the Alzheimer’s Disease Neuroimaging Initiative (ADNI) database. Our analysis focused on individuals with MCI who exhibit progressive deterioration in cognitive performance in comparison to those who remain stable over time (Fig. 1A). We thus considered data from two groups (Fig. 1B). First, subjects in the stable MCI (sMCI) group had a baseline diagnosis of MCI which was retained at follow-up, with at least 18 months between diagnoses. Secondly, subjects in the progressive MCI (pMCI) group were subjects who over the course of a similar duration progressed from a diagnosis of MCI to AD. To train our deep learning model (see below) we additionally analyzed data from CN subjects along with subjects with a diagnosis of AD. Two independent cohorts were analyzed for the two latter groups (data from ADNI-1 and ADNI-2/GO).

**Fig. 1.**
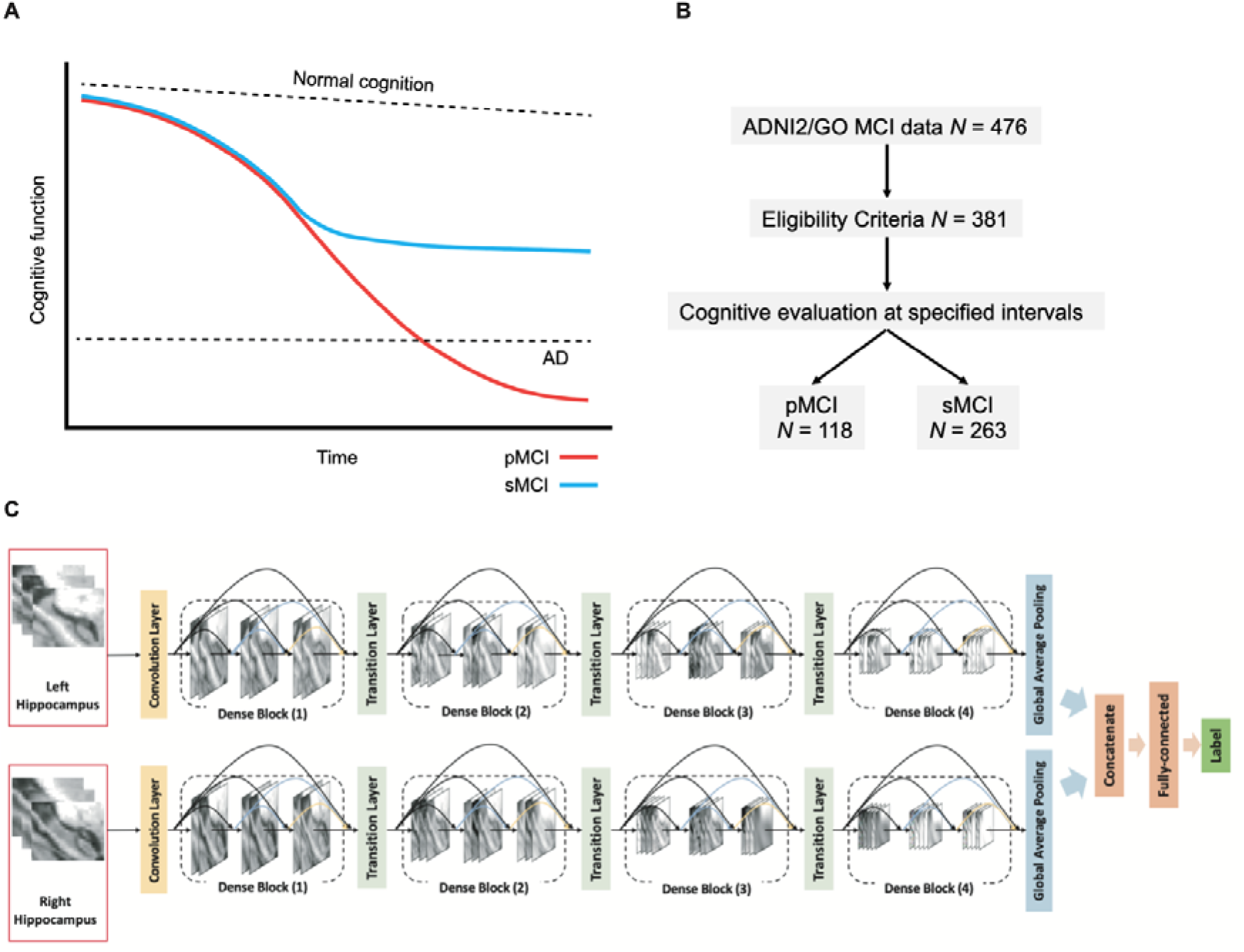
Study design and methods. (**A**) Hypothetical models of MCI progression. In pMCI gradual cognitive decline continues until individuals meet the diagnostic criteria of AD. In sMCI cognitive performance remains relatively stable over time. (**B**) Group allocation criteria. Cognitive evaluations at baseline and follow-up visits were used to classify subjects in the pMCI and sMCI groups. (**C**) Illustration of proposed deep learning model. Abbreviations: AD=Alzheimer’s disease, MCI=mild cognitive impairment, pMCI=progressive mild cognitive impairment, sMCI=stable mild cognitive impairment.

### Participant Characteristics

The demographic characteristics of subjects in the AD, CN, pMCI, and sMCI groups are shown in **Table 1**. Comparing the pMCI and sMCI groups, there were significant differences in age (*t_379_* = 2.449, *p* = 0.015) and MMSE score (Wilcoxon Rank-sum, *Z* = 4.938, *p* = 7.89 × 10^−7^). No significant difference was observed for gender distribution (χ^2^=0.072, *p* = 0.787) or education (*t_379_* = 0.756, *p* = 0.450). In the comparisons between the AD and CN groups, in both cohorts there were significant differences in education (ADNI −1: *t_425_* = 4.483, *p* = 9.47 × 10^−6^; ADNI-2/GO: *t_347_* = 2.614, *p* = 0.009) and MMSE score (ADNI-1: *Z* = 17.900, *p* = 1.19 × 10^−71^; ADNI-2/GO: *Z* = 15.953, *p* = 2.70 × 10^−57^). Age significantly differed in the ADNI-2/GO (*t_347_* = 1.901, *p* = 0.058) cohort, but not in ADNI-1(*t_425_* = 0.562, *p* = 0.575). Gender distributions were not significantly different in both cohorts (ADNI-1: χ^2^=0.022 *p* = 0.882; ADNI-2/GO: χ^2^=3.374, *p* = 0.066).

**Table 1.** Demographics.

|  | ADNI-1 |  | ADNI-2 & GO |  |  |  |
| --- | --- | --- | --- | --- | --- | --- |
|  | AD | CN | AD | CN | pMCI | sMCI |
| N | 197 | 230 | 159 | 190 | 118 | 263 |
| Age | 75.6 ± 7.7 | 76.0 ± 5.0 | 74.8 ± 8.1 | 73.4 ± 6.4 | 73.6 ± 7.1 | 71.7 ± 7.3 |
| Gender, female | 95 (48.2%) | 112 (48.7%) | 68 (42.8%) | 100 (52.6%) | 52 (44.1%) | 112 (42.6%) |
| Education | 14.7 ± 3.1 | 16.0 ± 2.8 | 15.8 ± 2.7 | 16.5 ± 2.6 | 16.0 ± 2.7 | 16.2 ± 2.7 |
| MMSE | 23.3 ± 2.0 | 29.1 ± 1.0 | 23.1 ± 2.1 | 29.0 ± 1.3 | 27.3 ± 1.8 | 28.3 ± 1.7 |

Continuous variables are presented as mean ± SD and categorical variable is presented as %. Abbreviations:
AD = Alzheimer's Disease, CN = Cognitively normal, pMCI = progressive mild cognitive impairment, sMCI = stable mild cognitive impairment, N = number of subjects, MMSE = Mini-Mental State Examination.

### A deep learning model for classifying stable and progressive MCI

We next developed a deep learning model, based on the DenseNet architecture (27) (Fig. 1C) for classification of pMCI vs. sMCI, as an initial step prior to delineating the role of hippocampal subfields in MCI progression. As input data, the model utilizes voxel intensity values from a 3D bounding box surrounding the entire hippocampal region. As in previous studies (16, 18) the model was trained to first differentiate AD and CN (data source: ADNI-2/GO), with the assumption that these 2 extreme classes would allow the model to learn the necessary representations for classifying MCI subjects as well. The model was then tested on the task of differentiating pMCI vs. sMCI (data source: ADNI-2/GO). We also validated the model’s performance by retesting it on the task of differentiating AD vs. CN (data source: ADNI-1). Data augmentation was applied within the training data set to improve the performance of the model and its generalizability. We used 10-fold cross-validation within the training set to optimize and fine-tune the model’s performance, finding similar performance across the different folds (*SI appendix*, Fig. S1). The model with the best performance achieved maximal accuracy of 94.07% in one of the folds, with an area under the curve (AUC) of the receiver operating characteristic (ROC) of 0.993. The model was further validated with AD and CN data from ADNI-1, achieving an accuracy of 86.20%, with an AUC of 0.937 (Fig. 2A). Thus, despite a decrease in accuracy, which may be expected given that ADNI-1 is based on lower MRI field strength (1.5T, relative to ADNI 2/GO’s 3T) the model was overall stable and robust. This model was thus used next for differentiating the pMCI and sMCI groups based on data from ADNI-2/GO. In this task, the model achieved an accuracy of 75.85% and an AUC of 0.777 (Fig. 2B). Repeating the analysis with an age-matched test cohort had minimal effects on the accuracy of the model (*SI appendix*, Fig. S2).

Overall our proposed deep learning-based classification framework corroborates earlier results, by demonstrating comparable accuracy performance to those reported previously for the classification of pMCI vs. sMCI (16–18). It establishes that whole hippocampus structural features can be used to differentiate pMCI from sMCI.

**Fig. 2.**
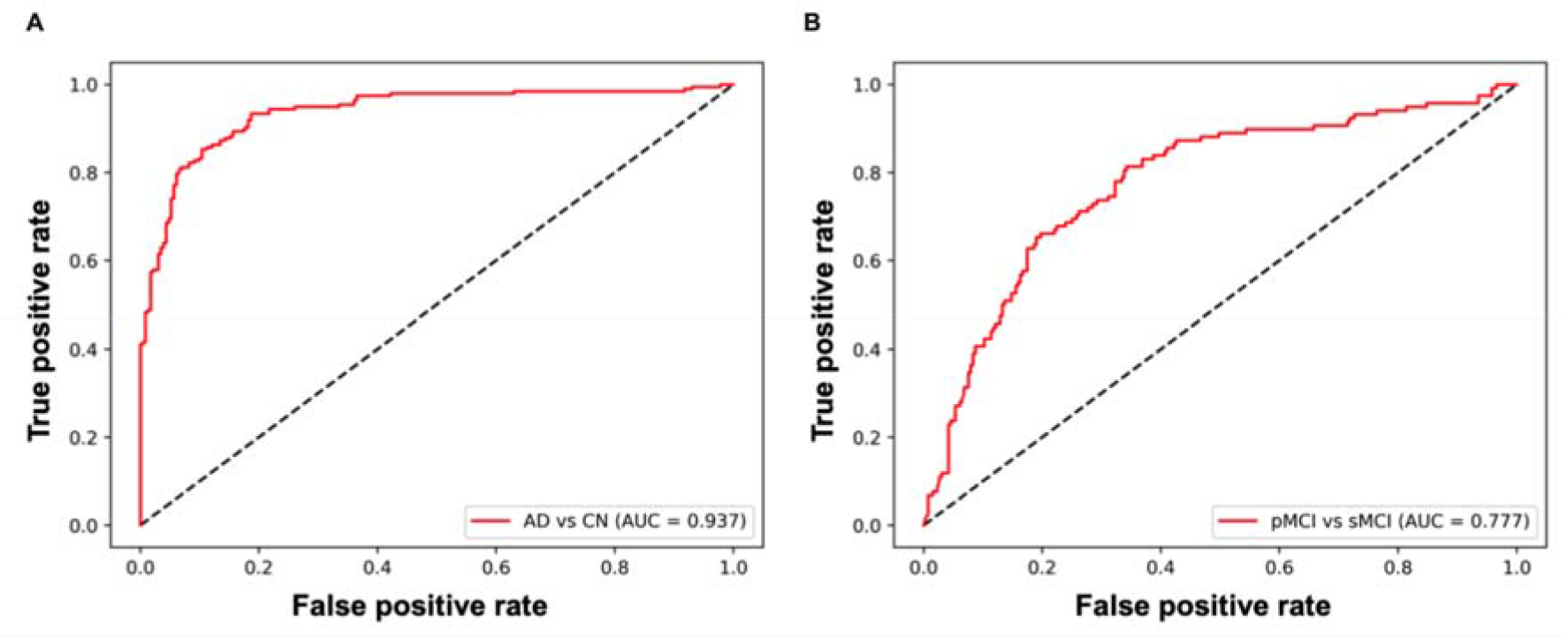
ROC curves showing the performance of the proposed deep learning model. (*A*) For the AD versus CN task. (*B*) For the pMCI vs sMCI task. Abbreviations: AD=Alzheimer’s disease, CN=cognitively normal, pMCI=progressive mild cognitive impairment, sMCI=stable mild cognitive impairment, ROC=receiver operating characteristic, AUC=area under the curve.

### Contribution of hippocampal subfields to MCI progression: occlusion analysis

We next sought to test if differentiation of pMCI from sMCI can be achieved with data derived from single hippocampal subfields, assessing the relative contribution of each subfield to classification performance. We first segmented the hippocampal subfields in each of the subjects using a validated automated method (28) (Fig. 3A). The contribution of each subfield was then assessed using an adaptation of occlusion analysis, a common approach in computer vision (e.g., (29)). Briefly, in this analysis we retested the deep learning model, each time occluding a bilateral binary mask of each of the hippocampal subfields from the model’s test data (i.e., from the 3D bounding box). The occlusion was achieved by setting the intensity values of each hippocampal subfield to zero in the input data. The performance (accuracy) of the models were then ranked and compared to each other as well as to the model based on an intact hippocampus (Fig. 3B). Accuracy levels for each of the models differed considerably (Fig. 3C).The occlusion of subfields led to a decrease in accuracy. In particular, occlusion of the subiculum, CA1, presubiculum, and molecular layer led to dramatic decreases in accuracy, relative to other subfields, including CA2/3 and CA4 for example. Thus, the presubiculum, subiculum, CA1, and the molecular layer had the largest impact on the performance of the model (Fig. 3D).

**Fig. 3.**
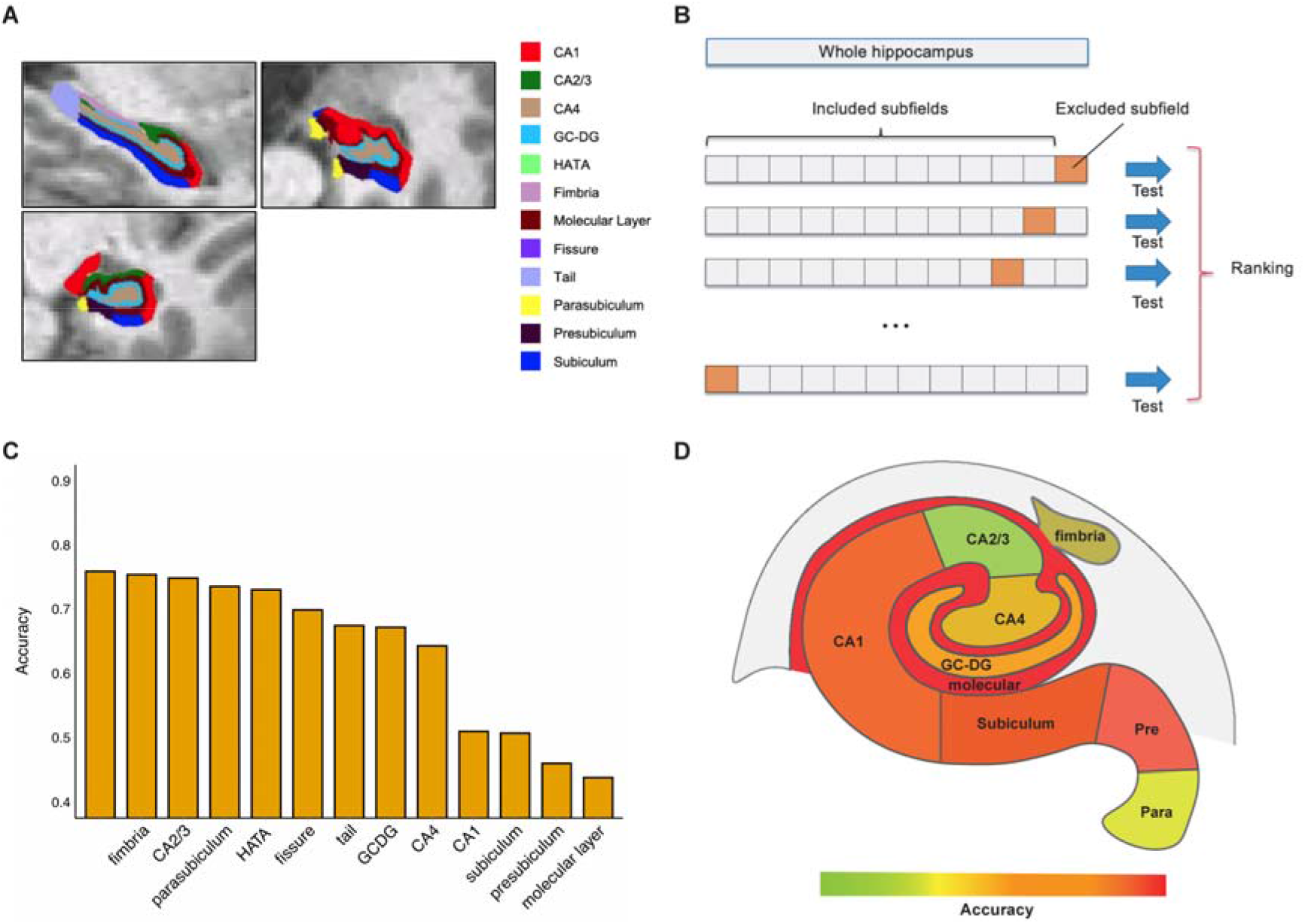
Occlusion analysis of hippocampal subfields. (*A*) Hippocampal subfield segmentation. The segmentation results of hippocampal subfields are illustrated on a single representative subject. (*B*) Schematic framework of the occlusion analysis. In each model, one of the hippocampal subfields was occluded (masked out) in the testing data and the performance of the model (its accuracy) was ranked relative to the occlusion of other subfields and to the performance of the original intact model. (*C*) The results of the occlusion analysis are shown for each model, along with the results of the original intact model (*D*) Accuracy performance of each model, superimposed on top of an illustration of the major hippocampal subfields (note: not all subfields are shown). Abbreviations: CA=cornu ammonis, HATA=hippocampus-amygdala-transition-area, GC-DG=granule cell layer of the dentate gyrus.

We also assessed the effect of laterality on the accuracy of classification by repeating the occlusion analysis with left vs. right hippocampal subfield masks tested separately. For the most impacted subfields (subiculum, CA1, presubiculum, and molecular layer), occlusion of the left hemisphere had a more pronounced effect on accuracy relative to occlusion of the right hemisphere (*SI appendix*, Fig. S3).

The occlusion analysis revealed that many of the subfields had little to no contribution to the performance of the model. Namely, occlusion of CA2/3 and parasubiculum, for example, resulted in an accuracy loss of less than 5%. This suggests that in the classification of pMCI and sMCI some of the subfields may be redundant. We evaluated this possibility by performing a sequential version of the occlusion analysis, where occlusion is accumulated from step to step (Fig. 4A), in descending order with respect to each subfield’s contribution to the model’s performance (as reported in Fig. 3). Accuracy was evaluated as a function of the ratio of the total occluded volume to the volume of the entire hippocampus. In comparison to the model with no occlusion, the accuracy started decreasing strongly when 10.5% of the volume of the hippocampus was removed (with occlusion of the fimbria, parasubiculum, and CA2/3), estimated by fitting the data with a log-sigmoid function, and extracting the resulting curve’s change points using a Bayesian inference approach (see Materials and Methods). In other words, we found that around 10.5% of the volume of the hippocampus was redundant in classifying pMCI versus sMCI (*SI appendix*, Fig. S4). Upon removal of more than 30.2% of the volume of the hippocampus accuracy levels started saturating (*SI appendix*, Fig. S4). Finally, repeating the occlusion analysis with age-matched data had little effect on the results (*SI appendix*, Fig. S5).

**Fig. 4.**
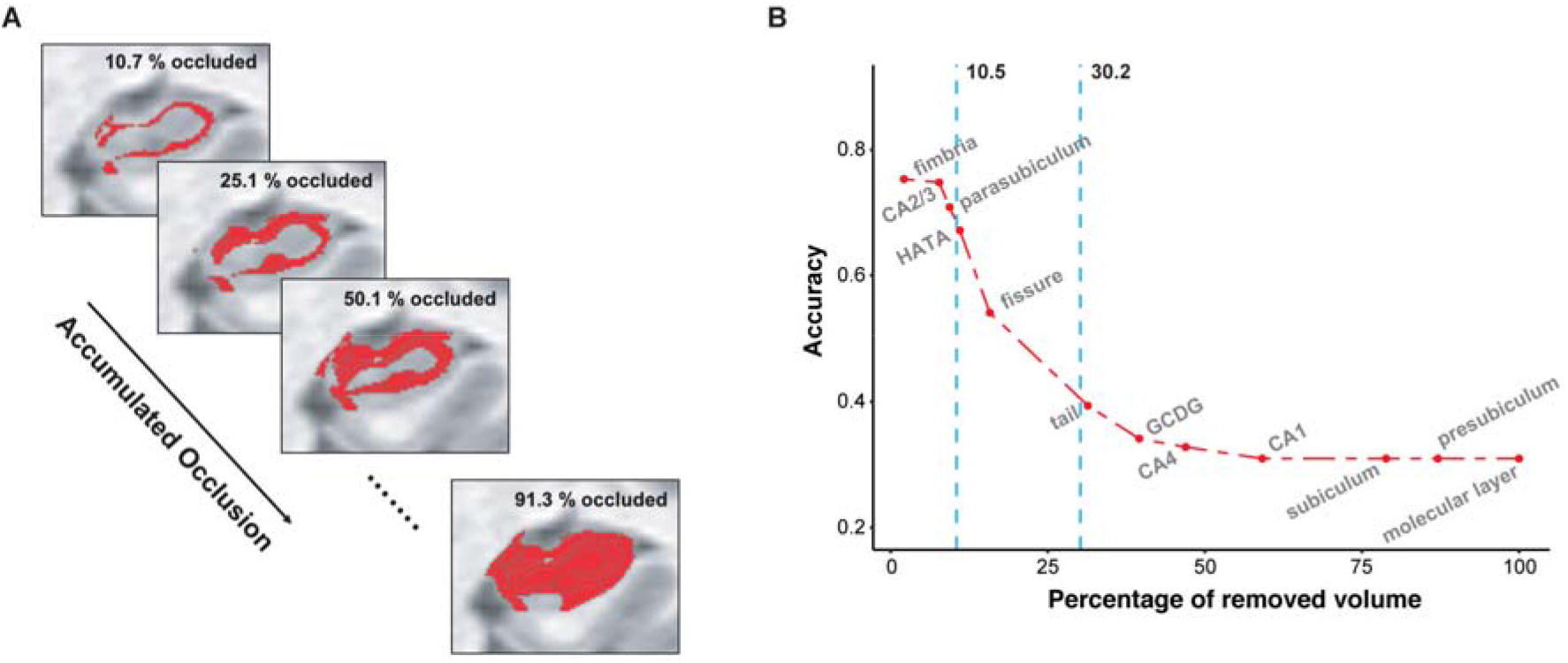
Accumulated occlusion analysis of hippocampal subfields. (*A*) An illustration of gradually accumulating occlusion. On each step, additional hippocampal volume is masked out of the analysis. (*B*) The model’s accuracy is shown as a function of accumulated occlusion, with subfields masked out in descending order according to their contribution to the model’s accuracy (as shown in Fig. 3C). The accuracy started decreasing strongly (i.e., relative to the accuracy of the full model) upon removal of 10.5% of the volume of the hippocampus, as identified with change-point analysis. Abbreviations: CA=cornu ammonis, HATA=hippocampus-amygdala-transition-area, GC-DG=granule cell layer of the dentate gyrus.

The occlusion of the presubiculum, subiculum, CA1, and molecular layer had the largest impact on the performance of the model. These 4 subfields are among the largest in the hippocampus, raising the possibility that their occlusion resulted in large losses in accuracy merely because of their large size. To test the possible confounding role of subfield size in our proposed framework, we repeated the occlusion analysis, with random occluded masks, which are of the same size as the presubiculum, subiculum, CA1, and molecular layer (Fig. 5A). We generated a null distribution for each of the 4 major subfields, based on 1000 random permutations, each of which consisting of random parcels, with the same size as the tested subfields but at random spatial locations (*SI appendix*, Fig. S6). The occlusion of the random parcels resulted in markedly lower loss of accuracy, relative to that observed in the original analysis (all *p*<0.001, Fig. 5B). This suggests that the loss of accuracy observed when the presubiculum, subiculum, CA1, and molecular layer were occluded was not merely a reflection of their size.

**Fig. 5.**
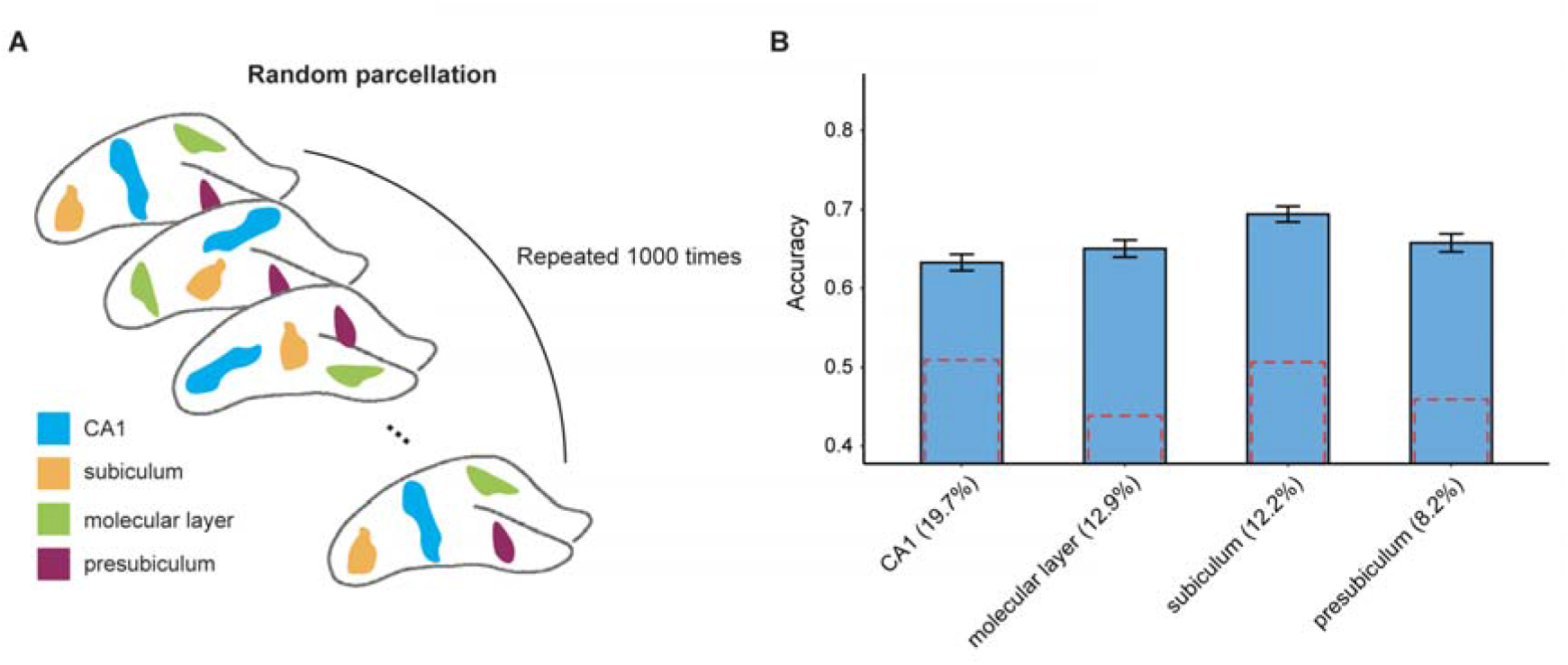
Random occlusion analysis for the major hippocampal subfields. (A) The potential confounding effect of subfield size was assessed by repeating the occlusion analysis 1000 times, each time occluding random parcels, which are of the same size as the presubiculum, subiculum, CA1, and molecular layer, but are placed at random spatial locations. (B) The results of the random occlusion analysis are shown with reference to each of the 4 hippocampal subfields (the relative size of each mask is indicated as % of total hippocampal volume). Dashed red lines denote the original accuracy loss observed (same as in Fig. 3C). Error bars denote standard deviation.

## Discussion

Individuals with MCI show strongly variable symptomatic trajectories, with some progressing eventually to a probable diagnosis of AD, while others showing a more stable pattern of cognitive performance over time. In this paper, we propose a novel framework for the analysis of the progression and stability in of MCI based on deep learning and occlusion analysis. First, we introduced a deep convolutional neural network model based on the DenseNet architecture (27) for classifying pMCI vs. sMCI. Second, we proposed a novel analytical framework based on occlusion analysis to evaluate the contribution of hippocampal subfields on the performance of the proposed deep learning model, thus assessing the role of the different subfields in the stability and progression of MCI. Finally, as a secondary step, we applied a gradually accumulating occlusion analysis which allowed us to assess the degree of redundancy in the hippocampal features in relation to the classification of pMCI and sMCI.

As an initial step prior to the evaluation of the role of hippocampal subfields, we trained a deep convolutional neural network to classify the pMCI and sMCI groups based on all structural hippocampus features. This model achieved an accuracy of 75.85% (and an AUC of 0.777). This classification performance in on par with earlier deep learning models developed to classify pMCI vs. sMCI based on whole hippocampus structural features or multi-model features (e.g., (30, 31)), which ranged from 72% to 76%. For example, a 3D-convolutional neural network based on multi-modal data, integrating structural MRI and positron emission tomography (PET) classified pMCI vs. sMCI with an accuracy of 72.22% (16), while another recently described hybrid convolutional and recurrent neural network based on internal and external hippocampal patches yielded classification accuracy of 72.50% for the same task (17). Another recent study utilized deep learning and hippocampal features predicting progression time from MCI to AD with a concordance index of 0.762 (18). Our model corroborates these earlier reports by demonstrating that prediction of MCI stability and progression can be achieved with good accuracy rates based solely on hippocampal features.

Our findings extend earlier reports by delineating the contribution of single hippocampal subfields to the progression and stability of MCI. Our implementation of occlusion analysis revealed marked differences between the subfields in differentiating the pMCI and sMCI groups. In particular, the results suggest that the subiculum, presubiculum, CA1, and molecular layer were more central to this classification task than any other subfield. While these are among the largest subfields in the hippocampus, our results also demonstrate that their contribution to MCI progression was not merely a reflection of their size. The findings are consistent with earlier reports on the involvement of CA1, the subiculum (24, 32), and the molecular layer (33, 34) in the progression of MCI. While several other studies implicated CA2/3, fimbria, and GC-DG in MCI progression (25, 26), our model suggests that these subfields play a more minor role. Our findings may reflect the neuropathological cascade characteristic of AD. Namely, neurofibrillary tangles in AD neurodegeneration progress from CA1 to the subiculum, before reaching CA2/3 (35), Neuronal loss in CA1 is prominent as AD neurodegeneration progresses, while being milder and slower in the subiculum (36). The results also reveal a more pronounced contribution of subfields in the left hemisphere to the performance of the model, consistent with earlier results (37, 38). Altogether, although our findings highlight the contribution of subiculum, presubiculum, CA1, and molecular layer in the progression of MCI further examination into the possible involvement of other subfields is warranted.

As a secondary step to the occlusion analysis, which helped us assess the contribution of single subfields to MCI progression and stability, we also evaluated how accumulated occlusion of hippocampal features affected the results. This analysis revealed that 10.5% of the volume of the hippocampus was redundant in the differentiation between pMCI and sMCI. These results may reflect the progressive nature of neurodegeneration in AD, wherein the redundant subfields have yet to have been affected, and thus do not yet differentiate among patients with stable and progressive disease trajectories. Another speculative possibility, which remains to be tested in future research, is that the redundancy reflects neuroprotective mechanisms that allow individuals with MCI to compensate for the earlier phases of neurodegeneration. Compensatory and reserve mechanisms have been widely postulated to operate in response to aging and neurodegeneration (39, 40). Future research could test if redundancy at the level of hippocampal structure is functionally advantageous, offering patients with a coping mechanism for early-phase neurodegeneration.

Several limitations should be noted. First, while our study considered longitudinal clinical evaluations we did not examine longitudinal changes in imaging metrics. Our focus here was on evaluations of prognostic markers of conversion from MCI to AD. Future research could use similar methods to examine longitudinal imaging data. Second, we considered a single imaging modality in our models (structural MRI). Studies have consistently revealed the superiority of multimodal features in diagnostic and prognostic models (e.g., (30)). Since our focus here was on hippocampal subfields, integration of data with lower spatial resolution like that obtained from PET Amyloid Imaging would have been challenging. Yet, we acknowledge that basing our models on a single modality may have reduced its performance in the classification task. Finally, it would be beneficial to replicate the results with data obtained with higher-resolution neuroimaging.

In conclusion, the current study delineates the contribution of hippocampal subfields to the progression and stability of MCI, highlighting the central role of the subiculum, presubiculum, CA1, and molecular layer in differentiation between pMCI and sMCI. The results further reveal that around 10.5% of the volume of the hippocampus is redundant in the differentiation between these two groups. These results highlight the need to consider the intricate structure of the hippocampus in studies of AD neurodegeneration.

## Materials and Methods

### Experimental data

Data used in the preparation of this article were obtained from the ADNI. The ADNI was launched in 2003 as a public-private partnership, led by Principal Investigator Michael W. Weiner, MD. The primary goal of ADNI has been to test whether serial MRI, other biological markers, and clinical and neuropsychological assessment can be combined to measure the progression of MCI and early AD. For up-to-date information, see www.adni-info.org. All subjects provided written informed consent and the study protocol was approved by the local Institutional Review Boards. Inclusion criteria for cognitively normal (CN) were as follows: (a) free of memory complaints. (b) Mini-Mental State Examination (MMSE) scores between 24-30. (c) Clinical dementia rating (CDR) score of 0. (d) Normal scores in the Logical Memory LJ subscale of the Wechsler Memory Scale-Revised, using education-adjusted cutoffs (41). Inclusion criteria for AD were as follows: (a) subjective memory concern (b) MMSE scores between 20-26. (c) CDR score of 0.5 or 1. (d) Abnormal scores in the Logical Memory LJ subscale of the Wechsler Memory Scale-Revised, using education-adjusted cutoffs. Inclusion criteria for MCI subjects were as follows: (a) subjective memory concern. (b) MMSE scores between 24-30. (C). Clinical dementia rating (CDR) score of 0.5. (d) Abnormal scores in the Logical Memory ◻ subscale of the Wechsler Memory Scale-Revised, using education-adjusted cutoffs. We used data from 349 subjects from ADNI-2/GO to train the deep learning model and evaluated the model with an independent cohort of 427 subjects from ADNI-1. An additional sample of 381 subjects with MCI at baseline, obtained from ADNI-2/GO, was classified as either sMCI or pMCI based on longitudinal diagnostic evaluations. We excluded subjects who were diagnosed with MCI at baseline but reverted to CN status during follow-up. The demographic characteristics of each cohort analyzed in this study are summarized in Table 1.

### Imaging data

Input data for the deep learning model were acquired at ADNI sites using 1.5T (ADNI-1) and 3T (ADNI-2/GO) scanners and were based on either an inversion recovery-fast spoiled gradient recalled (IR-SPGR) or a magnetization-prepared rapid gradient-echo (MP-RAGE) sequences (42). Full details of the image acquisition parameters are listed on the ADNI website (http://adni.loni.usc.edu/methods/documents/mri-protocols/).

### Image Processing

All images were corrected for intensity non-uniformity artifacts (43) and were registered into the MNI152 template using FMRIB Software Library v6.0 (44). Input data from the left and right hippocampus were extracted from the T1-weighted MRI images. We first defined a 3D bounding box of size 44 × 52 × 52 voxels around the hippocampal region. The bounding box’s size was sufficient in covering the hippocampal region in all our target subjects, with its size corresponding to that used in previous studies (17, 45). The voxel intensities within the bounding box were normalized into a range between 0 and 1. Intensity values from each voxel within the bounding box were then extracted and used as inputs in the deep learning model described below.

Data Augmentation: To artificially increase the size of the model’s training dataset, and improve its performance and generalizability, we used an image data augmentation technique with Scikit-learn 0.22.1 (46). The augmented image data were generated through the addition of noise with mean 0 and standard deviation 1, contrast enhancement by effectively spreading out the most frequent intensity values (stretching out the intensity range) and flipping left and right. In total, 954 AD images, and 1140 CN images were generated through augmentation and used in the training dataset to improve the performance of the model.

Hippocampal subfield segmentation: Subfields in the hippocampus were segmented with an automated segmentation tool available in FreeSurfer v6.0 (47), which is based on a new statistical atlas built primarily upon ultra-high resolution (~0.1mm isotropic), *ex vivo* MRI data. This approach uses Bayesian inference that relied on image intensities and a tetrahedral mesh-based probabilistic atlas of the hippocampal formation, constructed from a library of *in vivo* data and *ex vivo* labeled data (47, 48). This widely used method was validated with ADNI-based MCI and AD data (47), with its reliability established both within and across scanners (49). The left and right hippocampus were segmented into twelve subfields: CA1, CA2/3, CA4, hippocamps-amygdala transition area (HATA), granule cell layer of the dentate gyrus (GC-DG), fimbria, molecular layer, hippocampal fissure, hippocampal tail, subiculum, parasubiculum, and subiculum.

### Deep learning model architecture

A deep learning model based on the DenseNet architecture (27) was trained to learn relevant maps for classifying pMCI vs. sMCI. This state-of-the-art convolutional neural network architecture was chosen as it shows excellent classification performance with a range of datasets while diminishing the vanishing gradient problem and reducing the number of parameters (27). We used deep learning, rather than a more simple machine learning model, such as random forest, or support vector machine, primarily since similar models have achieved the best performance in previous studies (17, 18, 45), when based on complex features directly extracted from the raw data (i.e., whole hippocampal intensity values). Moreover, we assumed complex, possibly non-linear relationships between hippocampal subfields and the progression from MCI to AD, further motivating the use of deep learning.

The deep learning model (See Fig. 1C) comprised of two streams for the left and right hippocampus. Each stream consisted of a convolutional layer, 4 dense blocks, 3 transition layers, and a global average pooling layer. The outputs of the two streams were then concatenated as input to a fully connected layer. First, the image was passed through a stack of convolutional layers, where the filters were of size 5×5×5. The convolution stride was fixed to 1 voxel; The max-pooling layer had a stride of size 2×2×2 and a kernel size of 2×2×2. The dense block consisted of multiple convolution units, which were equipped with batch normalization layer, leaky rectified linear unit, a 1 × 1 × 1 convolutional layer, a 3×3×3 convolutional layer and a dropout layer. Every convolutional unit was connected to all previous layers by shortcut connections. A transition layer allowed for dimensionality reduction of feature maps in between dense blocks. It was composed of a batch normalization layer, leaky rectified linear unit, a 1×1×1 convolutional layer, a 3×3×3 convolutional layer, a dropout layer and an averaging pooling layer. The stacks of global averaging pooling layers were concatenated and connected by a fully-connected layer. The output value was processed by the fully-connected layer with a sigmoid activation function.

### Implementation

The deep learning model was built with the Keras application programming interface in TensorFlow 2.0. Training and testing of the model were carried out with an Ubuntu 18.04.3 operating system and two Nvidia Tesla V100 graphic cards with 16GB memory each. The model was parallelized across graphic cards. We trained the model with a mini-batch size of 64 and 200 epochs. The deep learning model was optimized using stochastic gradient descent (50) with momentums and an exponentially decaying learning rate. The initial learning rate was 0.0001 and it was decayed by 0.9 after every 10000 steps. We added a dropout layer in the dense block and set the dropout rate to 0.2. In the batch normalization, beta and gamma weight were initialized with L2 regularization set at 1×10^−4^ and epsilon set to 1.1 × 10^−5^. The L2 regularization penalty coefficient was set at 0.01 for the fully connected layer. The deep learning model was stable after an iteration of 150 epochs. We used a categorical cross-entropy loss function in our model. A focal loss function based on cross-entropy loss for imbalanced data was also used, but it had no differential effect on the performance of the model.

### Validation framework

We evaluated the performance of the proposed deep learning model using k-fold cross-validation, which allowed for the optimization of hyperparameters in the train set. We thus split the entire ADNI-2/GO data into k folds, selecting the parameters with the best performance and then applied the selected parameter to the test data to validate the performance of the proposed model.

### Implementation of occlusion analysis

Occlusion analysis was used for investigating the contribution of each hippocampal subfield to the performance of the prediction model. Masks for each hippocampal subfield were generated based on the subfield segmentation procedure described above (See Image Processing). We then masked out each hippocampal subfield (setting voxels of each hippocampal subfield to zero) from the input data of the test phase and retested the trained deep learning model as described before. We occluded left and right subfields simultaneously. Each occluded hippocampal subfield was thus ranked based on the performance (accuracy) of the model, relative to the original model, where input data from the entire hippocampus was used.

In a second implementation of occlusion analysis, we evaluated the performance of the prediction model under gradually accumulating occlusion of hippocampal subfields. This was achieved by retesting the prediction model, each time masking out an additional subfield (i.e., starting from one masked out subfield, then two etc.). Subfields were masked out sequentially, in descending order, according to their contribution to accuracy (as identified in the initial occlusion analysis). This step also allowed us to evaluate the stability of the occlusion analysis, ensuring that did not result in abrupt changes in classification performance. We additionally estimated the points in which the accumulated occlusion analysis showed large changes in accuracy. First, the results of the analysis (i.e., changes in accuracy observed in each of its 12 steps) were fitted with a nonlinear log-sigmoid curve (51). We then extracted the fitted curve’s change points using the R mcp package (52) by estimating change points between pre-defined models based on a Bayesian inference approach. In this approach change points correspond to the points in the curve where the data’s best predictive model changes from one model to another.

Lastly, we evaluated the potential confounding effect that the size of the occluded subfields had on the accuracy of classification through random occlusion analysis. This post-hoc analysis focused on the 4 major subfields which showed the most prominent effects in the original occlusion analysis (presubiculum, subiculum, CA1, and molecular layer). The average size (volume) of each of these 4 subfields was first calculated. We then generated, for each subfield, random parcels with identical size, but at random spatial locations based on a region growing algorithm (53). A total of 1000 random parcels were generated for each subfield. We then repeated the occlusion analysis 1000 times, occluding the random parcels at each permutation. This allowed us to generate a null distribution of accuracy values calculated based on the randomly placed masks. Differences between the original accuracy values and the values obtained via random occlusion were then assessed with a two-sided permutation test.

## Supporting information

Kwak_SI

## Author contribution

K.K and E.D conceived research; K.K. analyzed data; K.K, M.N, K.S.G, M.S and E.D interpreted results. K.K and E.D. wrote the paper.

