## Supplementary material for "The contribution of hippocampal subfields to the progression of neurodegeneration": Kwak_SI

^†^Data used in the preparation of this article were obtained from the Alzheimer's Disease Neuroimaging Initiative (ADNI) database ([http://adni.loni.usc.edu](http://adni.loni.usc.edu/)). As such, the investigators within the ADNI contributed to the design and implementation of the ADNI and/or provided data but did not participate in analysis or writing of this article. A complete listing of ADNI investigators can be found at <http://adni.loni.usc.edu/wp-content/uploads/how_to_apply/ADNI_Acknowledgement_List.pdf>.

This PDF file includes:

Figures S1 to S6

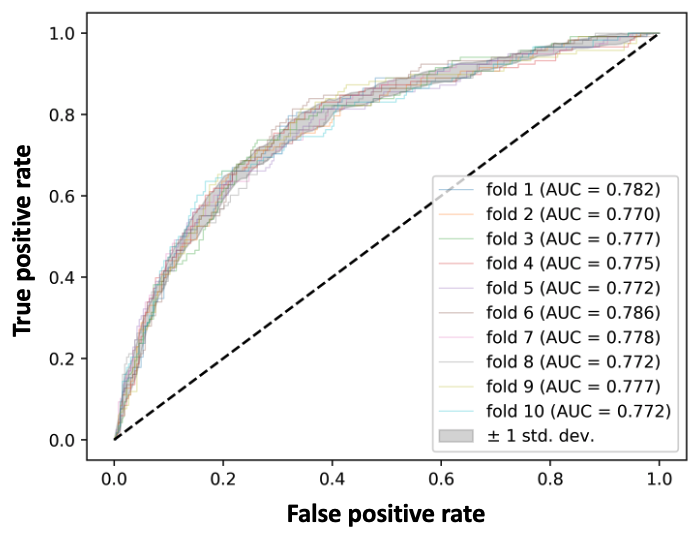

**Fig. S1. ROC curves for the proposed deep learning model obtained in each cross-validation fold.** ROC analysis revealed consistent performance of the proposed model during the training phase. Abbreviations: ROC=receiver operating characteristic, pMCI=progressive mild cognitive impairment, sMCI=stable mild cognitive impairment, AUC=area under the curve.

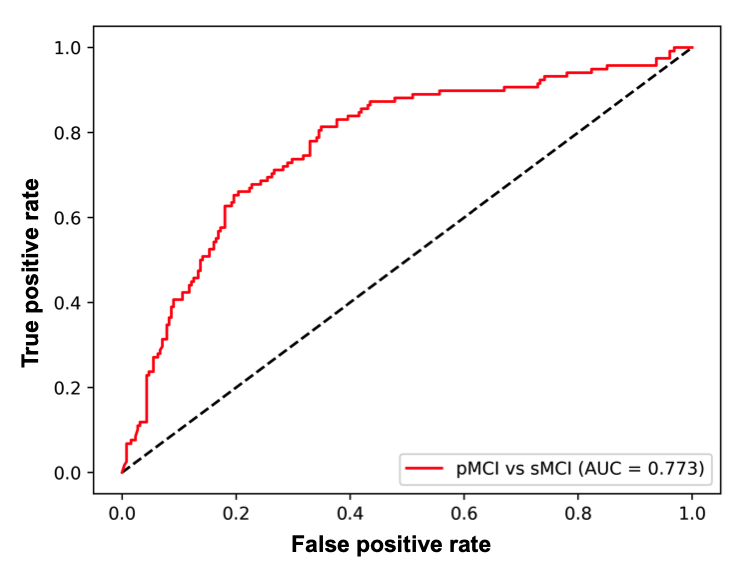

**Fig. S2. ROC curve of the proposed deep learning model for age-matched data.** The model was retested with age-matched subjects in the pMCI and sMCI groups (Age= 73.6$\pm$7.1; N= 118 and Age= 72.13$\pm$6.9; N=255 in the pMCI and sMCI groups, respectively). Age-matching was achieved by removing the minimal number of subjects required in order to eliminate the significant age differences between the groups. The model yielded an accuracy of 75.33% and an AUC of 0.773. Abbreviations: ROC=receiver operating characteristic, pMCI=progressive mild cognitive impairment, sMCI=stable mild cognitive impairment, AUC=area under the curve.

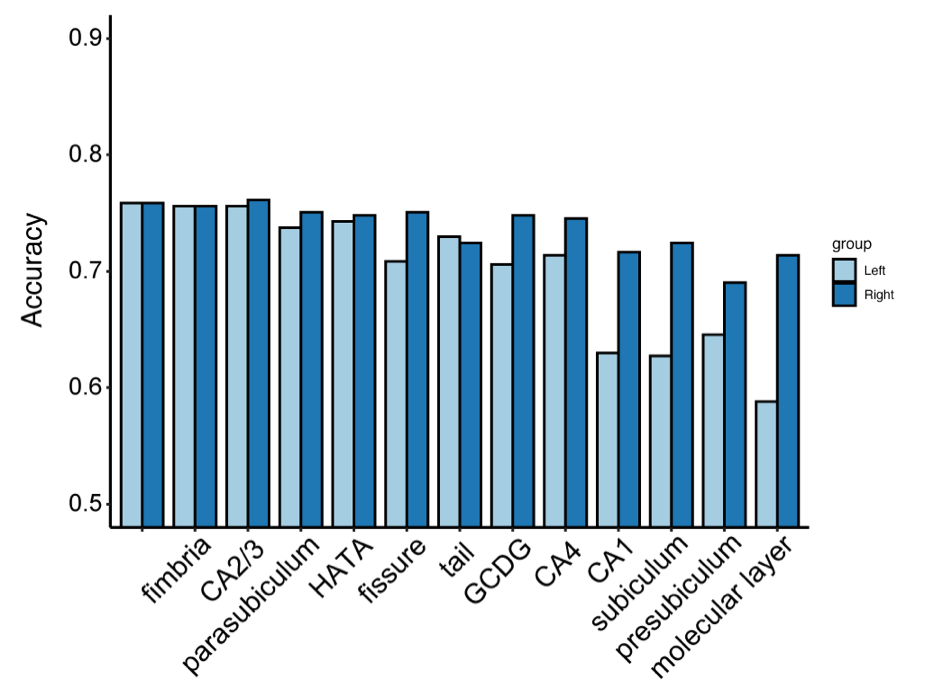

**Fig. S3. Effect of laterality in the occlusion analysis.** The occlusion analysis was repeated, with left vs. right masks occluded separately. Asymmetry was observed for CA1, subiculum, presubiculum and molecular layer where occlusion of the left hemisphere had a more pronounced effect than occlusion of the right hemisphere. Abbreviations: CA=cornu ammonis, HATA=hippocampus-amygdala-transition-area, GC-DG=granule cell layer of dentate gyrus.

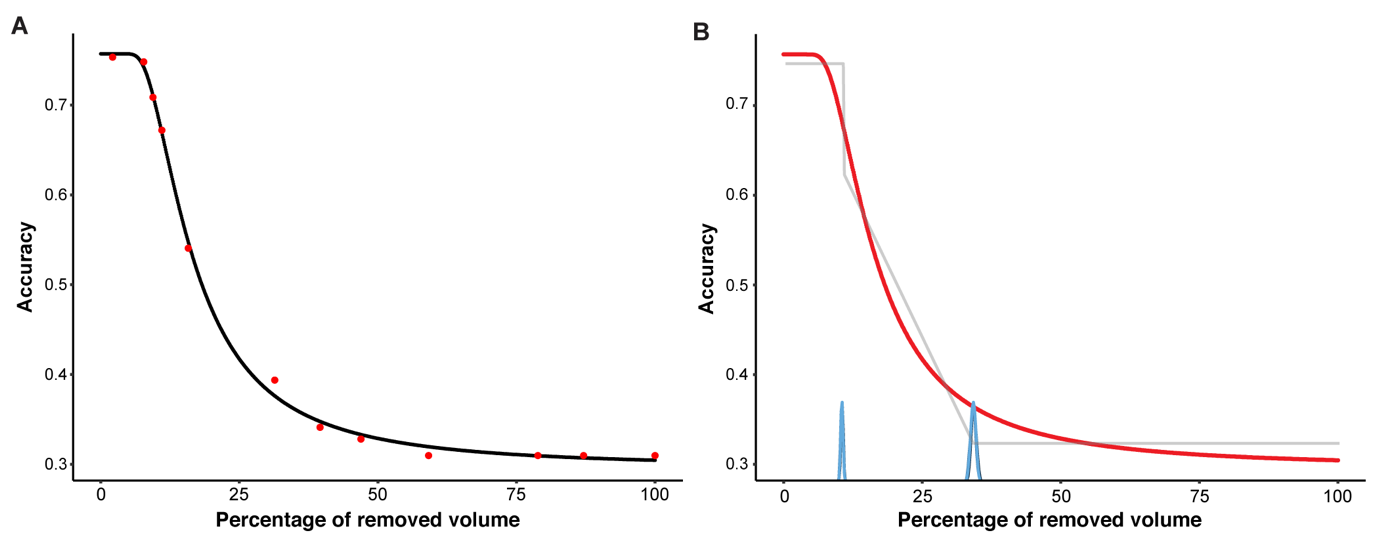

**Fig. S4. Change point detection in the accumulated occlusion analysis.** (*A*) A nonlinear log-sigmoid curve was fitted to the results of the accumulated occlusion analysis (shown is the curve with the best fit; r = 0.99). (*B*) The fitted curve (red line) with posterior lines (gray lines) are shown. The blue lines correspond to the posterior density of the curve’s change points.

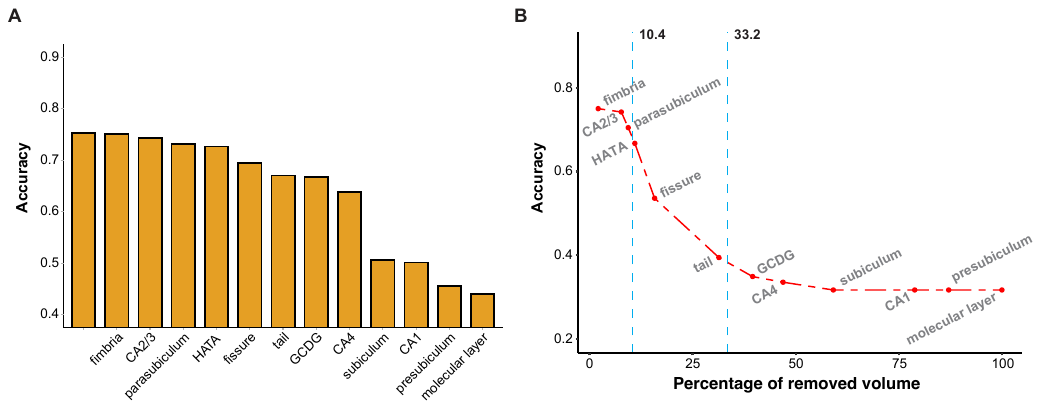

**Fig. S5. Results of the occlusion and accumulated occlusion analysis with age-matched data.** (*A*) The results of the occlusion analysis are shown. Using age-matched data had little effect on the results (maximal % change in accuracy = 0.762). (*B*) The model's accuracy is shown as a function of accumulated occlusion. The accuracy started decreasing upon removal of more than 10.4% of the volume of the hippocampus, as in the full model. Abbreviations: CA=cornu ammonis, HATA=hippocampus-amygdala-transition-area, GC-DG=granule cell layer of dentate gyrus.

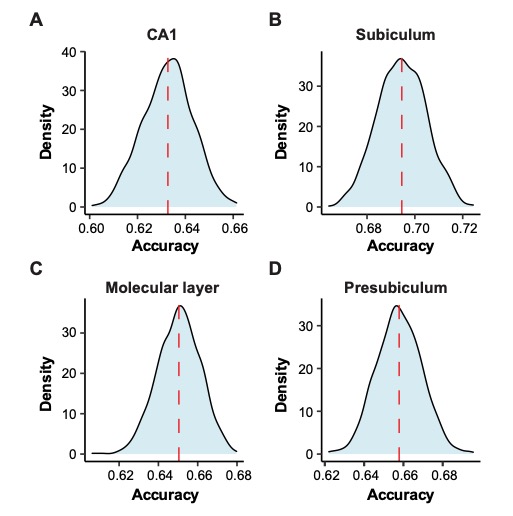

**Fig. S6. Null models obtained in the random occlusion analysis.** The distribution of accuracy values achieved in the random occlusion analyses are shown with respect to each subfield size. Red dotted lines denote the mean accuracy. Results are shown for random parcels corresponding to the size of: (*A*) CA1. (*B*) subiculum. (*C*) molecular layer. (*D*) presubiculum.
